# Genetic and Environmental interactions contribute to immune variation in rewilded mice

**DOI:** 10.1101/2023.03.17.533121

**Authors:** Oyebola Oyesola, Alexander E. Downie, Nina Howard, Ramya S. Barre, Kasalina Kiwanuka, Kimberly Zaldana, Ying-Han Chen, Arthur Menezes, Soo Ching Lee, Joseph Devlin, Octavio Mondragón-Palomino, Camila Oliveira Silva Souza, Christin Herrmann, Sergei Koralov, Ken Cadwell, Andrea L. Graham, P’ng Loke

## Abstract

The relative and synergistic contributions of genetics and environment to inter-individual immune response variation remain unclear, despite its implications for understanding both evolutionary biology and medicine. Here, we quantify interactive effects of genotype and environment on immune traits by investigating three inbred mouse strains rewilded in an outdoor enclosure and infected with the parasite, *Trichuris muris*. Whereas cytokine response heterogeneity was primarily driven by genotype, cellular composition heterogeneity was shaped by interactions between genotype and environment. Notably, genetic differences under laboratory conditions can be decreased following rewilding, and variation in T cell markers are more driven by genetics, whereas B cell markers are driven more by environment. Importantly, variation in worm burden is associated with measures of immune variation, as well as genetics and environment. These results indicate that nonheritable influences interact with genetic factors to shape immune variation, with synergistic impacts on the deployment and evolution of defense mechanisms.

## INTRODUCTION

An individual’s immune phenotype is shaped by some combination of genetic factors and nonheritable influences such as environmental exposure (including infection history and the microbiome) (1–9). However, the relative and potentially interacting contributions of heritable and nonheritable factors to inter-individual immune variation remain controversial despite the importance of such variation for both medicine and evolutionary biology. For example, variation in immune responses can determine whether an individual will experience severe or asymptomatic infection (10, 11), and whether severity arises due to failure to control pathogens or excessive collateral tissue damage following defective immune regulation (3).

Recent studies on the human immune system have aimed at identifying the relative contributions of genetic and environmental factors to variation in immune phenotypes among healthy individuals (1, 6, 7), as well as during infection (7) or inflammatory conditions (12). Such studies draw upon analysis of immunological divergence between identical twins (7) or genetic heritability estimation for immune traits through functional genomics (1). However, the design of these studies often makes the interactive effects of genetics and environment challenging to quantify(13, 14) and generally, has not been the well-examined in most immunological studies. For example, variation not attributable to genetics is generally attributed to environment alone rather than the possibility of Genotype-by-Environment interactions (G*E), which are inferred if effects of environment are differentially amplified in different genotypes, or vice-versa (15, 16). Important context-dependency in immune function (17) is thus often missing from these calculations. For example, what if the impact of environment upon memory T cell frequencies depends upon host genotype, or if, conversely, the impact of genotype upon memory T cell frequencies depends upon environment? Evolutionary biology is explicitly interested in such context dependencies because they provide the raw materials for adaptive evolution and diversification, indeed, genotype-by-environment interactions are common and substantial in effect for a variety of traits (18, 19) and disease outcomes (15, 16, 20). Here, we use mouse experiments together with statistical frameworks that quantify interactions in addition to the independent (or main) effects of genetics and environment to elucidate causes of variation in experimental immunology.

Controlled experiments with mice could help decipher interactions between genetic and environmental effects on the immune system, but most studies in mice instead aim to reduce environmental variation to discover genetic factors regulating cellular and molecular components of immunity (3, 21–23). Most times, this approach ignores interactions and provides only partial insight into direct genetic effects by ignoring the extent of the measured genetic effect that is mediated by the particular environment. We have taken a decidedly different approach of using an outdoor enclosure system to introduce laboratory mice of different genotypes into a natural environment that we term “rewilding” (24). In this particular study year, we tracked behavior outdoors(25), recover the mice for analysis, and then investigate genetic and environmental contributions to immune phenotypes (23, 24, 26). Previously, using mice with mutant alleles in inflammatory bowel disease susceptibility genes (*Nod2* and *Atg16l1*), we found that the genetic mutations affected the production of cytokines in response to microbial stimulation, whereas immune cell composition was more influenced by environment (23, 26). We also found that rewilded C57BL/6 mice become more susceptible to infection with the intestinal nematode parasite *Trichuris muris* (24). However, those experiments explored limited genetic variation and did not examine whether *interactions* between genetics and environment would influence immune phenotype and helminth susceptibility (4, 27).

Here, to quantify relative and interactive contributions of genetic and environmental influences on heterogeneity in immune profiles and helminth susceptibility, we compared C57BL/6, 129S1, and PWK/PhJ mice kept in a conventional vivarium versus those that were rewilded and measured inter-individual heterogeneity in immune cellular composition, cytokine responses and parasite burdens. Single cell RNA sequencing (scRNAseq) from individual mice also provided both cellular and functional readouts, which was associated with susceptibility to parasite infection. Together, our results demonstrate that interactions between genetics and environment are an important source of variation for specific immune traits, but there are also tissue dependent differential effects of environment versus genetics on specific cellular compartments such as T cells and B cells. Hence, we argue that our rewilding approach can demonstrate that the effect of an extreme environmental shift on immune phenotype is modulated by genetics, and the genetic effect is modulated by an environment. Such interactions are an important source of inter-individual immune variation.

## RESULTS

### Experimental Design

To quantify sources of heterogeneity in immune profiles and helminth susceptibility, we compared 3 different strains of mice - C57BL/6, 129S1, and PWK/PhJ mice – housed in two different environments - a conventional vivarium but kept in summerlike temperatures and photoperiods (hereafter “Lab” controls) versus those that were outdoor (hereafter “rewilded”) (Fig 1A). These host strains differ by up to 50 million single nucleotide polymorphisms (SNPs) and short insertions/deletions (indels) (*28, 29*); for perspective on this range of variation, the entire human population is estimated to contain approximately 90 million SNPs and indels (*30*). We rewilded mice (n = 72) or kept them in laboratory housing (n = 63) for 2 weeks and then infected them with approximately 200 eggs of the intestinal helminth *Trichuris muris* (n = 61) or left them uninfected (n = 74), returning them to the outdoor or vivarium environment for a further 3 weeks. We conducted two replicate experiments across different periods during the summer months (Block 1, n = 61, ending in July; Block 2, n = 74, ending in August). We collected blood and mesenteric lymph nodes (MLN) for immune phenotyping and fecal samples for microbiota analyses. To assess immune cell composition, we analyzed complete blood counts with differential (CBC/DIFF) of total blood, peripheral blood mononuclear cells (PBMCs) by flow cytometry with a lymphocyte panel (Table S1) and MLN cells with both a lymphocyte and myeloid cell panel (Table S1 and S2). To assess cytokine responses, we measured plasma cytokine concentrations and stimulated MLN cells with microbial antigens and measured cytokines released in the supernatant. Single cell RNA sequencing (scRNAseq) of MLN cells enabled phenotyping of both immune cell composition and function. We also assessed worm burden and worm prevalence in all mice exposed to *T. muris* at day 19-21 post infection prior to full worm maturation, which was necessary to prevent shedding of *T. muris* eggs into the rewilded environment. Serology and PCR screening panels tested for over 30 pathogens indicated that the mice had no other detectable infections (Table S3 and S4).

**Fig. 1:**
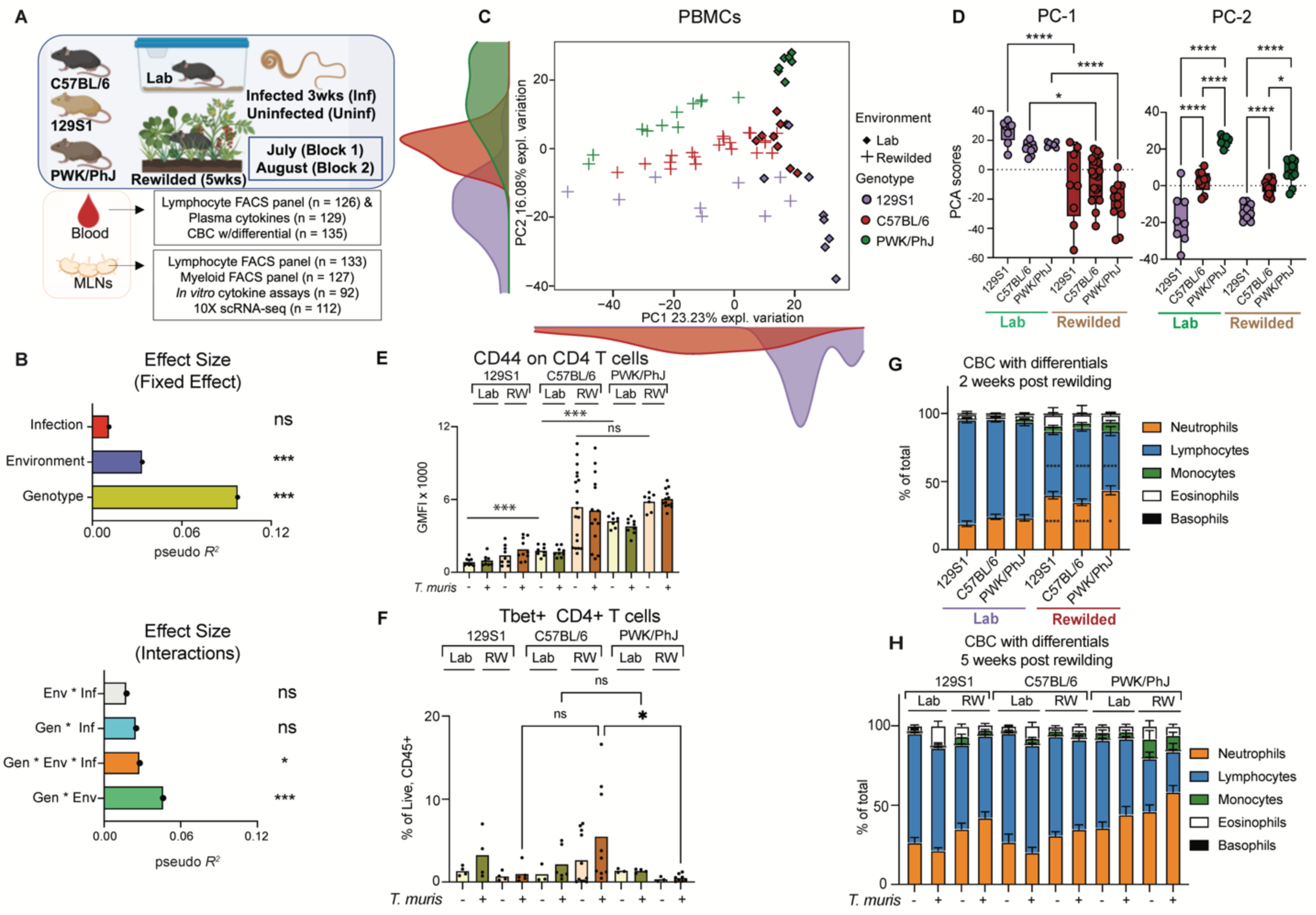
Interactions between Genotype and Environment contribute to variation in immune composition in murine PBMCs. (**A**) 51 C57BL/6, 41 129S1, 43 PWK/PhJ different mice strains (total =135) were used in these experiments. Some were kept in conventional vivarium (n = 63), and some were housed in the wild enclosure (Rewilded), (n = 72) for total of 5 weeks. In addition, some were exposed to 200 *T. muris* L3 eggs (n =61) while the others were left unexposed (n = 74). Experiments was repeated twice in July, Block 1 (n = 61) and August, Block 2 (n = 74). Blood was collected for Flow cytometry, Plasma cytokine assessment and CBC analysis. MLN cells were also collected for Single cell RNA sequencing (ScRNA-seq), Flow cytometry with two different panels - lymphocyte and myeloid panel as well as cytokine profiling of supernatants from MLN stimulated cells. For Flow cytometry visualization, only Block 2 was used for downstream analysis. (**B**) Bar plots showing the pseudo *R^2^* measure of effect size of predictor variables and interactions as calculated by multivariate distance matrix regression analysis (MDMR) (**C**) PCA of immune cell clusters identified by unsupervised clustering in the blood and the density of each population along the PCA and (**D**) Bar plot showing variance on PC1 and PC2 axis of PCA plots in (B). (**E**) Barplot showing GMFI of CD44 on blood CD4+ T cells, (**F**) % of Tbet+ CD4 T cells of Live, CD45+ T cells. Bar plots showing percentage of Neutrophils, Lymphocytes, Monocytes, Eosinophils, and Basophils out of total at (**G**) 2 weeks post re-wilding and (**H**) 5 weeks post re-wilding based on assessment by CBC with differentials. Statistical significance was determined by a based on MDMR analysis with R package (**B**) or based on one-way ANOVA test between different groups with Graph-Pad Software (**D**), (**E**) and (**F**). For (**E**) and (**F**) direct comparison was done between groups of interest with one-way ANOVA test. Data are displayed as mean ± SEM. *ns p>0.05; * p<0.05; ** p<0.01; *** p<0.001; **** p<0.0001*

### Interactions between Genotype, Environment and Infection contribute to variation in immune composition in murine PBMCs

Since most human immunology studies are conducted by analysis of PBMCs, first, we used this same approach to assess the immune phenotype of laboratory and rewilded mice. The immune cell composition of PBMCs in the peripheral blood were analyzed by spectral cytometry with a lymphocyte panel (Table S1). To quantify the relative contributions of Genotype (i.e. strain), Environment (i.e. Lab vs Rewilded) and Infection (i.e. exposure to *Trichuris muris*) and their interactions to the high-dimensional spectral cytometry data from the PBMC analysis, we used Multivariate Distance Matrix Regression (MDMR) analysis, a statistical approach used to determine factors contributing to variations for high-dimensional data (31–33). The MDMR model we used incorporated Genotype, Environment and Infection as fixed effects and the two independent experiments in July or August (Block) as a random effect to calculate the interactive and independent contribution of these factors to the outcomes. The MDMR model calculates the effect size of each variable on the outcome measure (here, cellular composition based on spectral cytometry data) to generate a pseudo-*R*^2^ value that quantifies the effect of the predictor variable on the dissociated outcome variable. The cellular composition data for the PBMCs of each individual mouse is determined by unsupervised K means clustering to group cells into clusters based on similarities of cellular parameters (Fig S1A). We calculate the composition of cells for each individual sample based on cluster membership established by K means clustering and this unbiased cluster composition data (Fig S1B) is then used as the outcome variable for the MDMR analysis.

The results from the MDMR analysis on PBMC cellular composition (Table S5) showed that Genotype and Environment had a significant effect on variation in cellular composition, not only as independent variables (Fig. 1B top) but also through interactions between Genotype and the Environment (Gen*Env) (Fig. 1B bottom). These patterns can be visualized through a principal component analysis (PCA) on cellular composition data of individual mice (Fig. 1C, 1D, Fig. S1B and Fig. S1C). The PCA plot indicated strong effects of environment on variation along the PC1 axis (Fig. 1C. and Fig. 1D) and of genetics on variation along the PC2 axis (Fig. 1C. and Fig. 1D), while infection displayed minimal effect (Fig. 1B, Fig. S1D and Fig. S1E). Variation along the PC1 axis for rewilded mice is substantially greater than for lab mice (Fig. 1C and 1D). The PCA plot also suggested that variance on the PC2 axis between mouse strains was greater in laboratory mice than for rewilded mice (Fig. 1C and 1D).

The loading factors in the PCA analysis showed that variation in expression of CD44 on CD4+ T cells is important for driving the differences that defines genetic variation on the PC2 axis (SFig. 1C). Although there are substantial difference in expression of CD44 on CD4+ T cells between the in-bred strains for lab-housed mice, these differences were no longer present between the C57BL/6 mice and the PWK/PhJ mice following rewilding (Fig 1E). Hence, the genetic differences seen in the clean laboratory environment can be reduced following rewilding. In contrast, rewilded C57BL/6 mice had more CD4^+^Tbet^+^ cells after infection compared to the PWK/PhJ mice, however, in laboratory conditions, there is no difference between these two strains of mice (Fig. 1F). These results indicate that a stronger T_H_1 response to *T. muris* in the C57BL/6 mice is only observed in the rewilding condition. Hence, genetic differences in response to infection can sometimes emerge only in rewilding conditions. These results illustrate how Gen*Env and Gen*Env*Inf interactions affect specific immune traits.

Complete blood count with differential (CBC/DIFF) is a standard clinical test used to assess inflammatory responses in patients. CBC/DIFF of total blood cells collected at 2 weeks and 5 weeks post rewilding provided data (Table. S6) on longitudinal changes in immune cell composition during rewilding, allowing us to compare the acute effects of environmental change (at 2 weeks post rewilding) to when the immune system has acclimatized to the new environment (at 5 weeks post rewilding). At 2 weeks post rewilding, there is a significant effect of rewilding on circulating neutrophils, lymphocytes and eosinophils across all genotypes (Fig. 1G). However, by 5 weeks post rewilding, we a trend towards more neutrophils in the rewilded PWK/PhJ mice than the other strains (Fig. 1H), indicating that this genotype might have a larger effect on neutrophil abundance especially in the rewilding environment. In contrast, eosinophils are more readily induced by *T. muris* infection in the laboratory mice at 5 weeks (Fig. 1H), than in the rewilded mice on the 129S1 and C57BL/6 background at 5 weeks. Together, these results indicated that acute environmental change (at 2 weeks) had a bigger effect on total blood cell composition than at 5 weeks. Additionally, infection induced responses in the laboratory setting can be altered during rewilding in specific genotypes.

Overall, these results suggest that there are important context dependent effects of genotype, environment, infection depending on the immune traits that are being measured in the peripheral blood and the timing of the environmental change.

### Interactions between Genotype and Environment contribute to variation in immune composition in murine secondary Mesenteric Lymphoid Organ

In contrast to what is obtainable in most human studies, murine studies give us an opportunity to easily assess immune responses in secondary lymphoid organs such as the lymph nodes. Therefore, we analyzed the mesenteric lymph nodes (MLNs) because they drain the intestinal tissues, which are most affected by *T. muris* infection (in the cecum), as well as by alterations to the gut microbiota. In contrast with the blood, MDMR analysis of immune composition of the draining MLNs based on unsupervised clustering of cells, as explained above with the lymphoid panel (Table S7), showed a significant effect of Genotype, Environment and Exposure to *T. muris* (Infection) in determining immune variation (Fig. 2A, left). Critically, an interactive effect of Genotype, Environment and Infection (Gen*Env*Inf) also contributed to the variation in immune composition in the MLN (Fig. 2A, right). We could visualize the contribution of Genotype, Environment and Infection with *T. muris* to MLN immune composition through the PCA analysis of immune cellular compositional data from individual mice (Fig. 2B and SFig. 2A). The PCA plot showed prominent effects of Genotype on variation along the PC1 axes (Fig. 2B and Fig. 2C) with effects of Environment along the PC2 axes (Fig. 2B and Fig. 2C) and *T. muris* Infection along the PC3 axes (Fig. S2B and S2C).

**Fig. 2:**
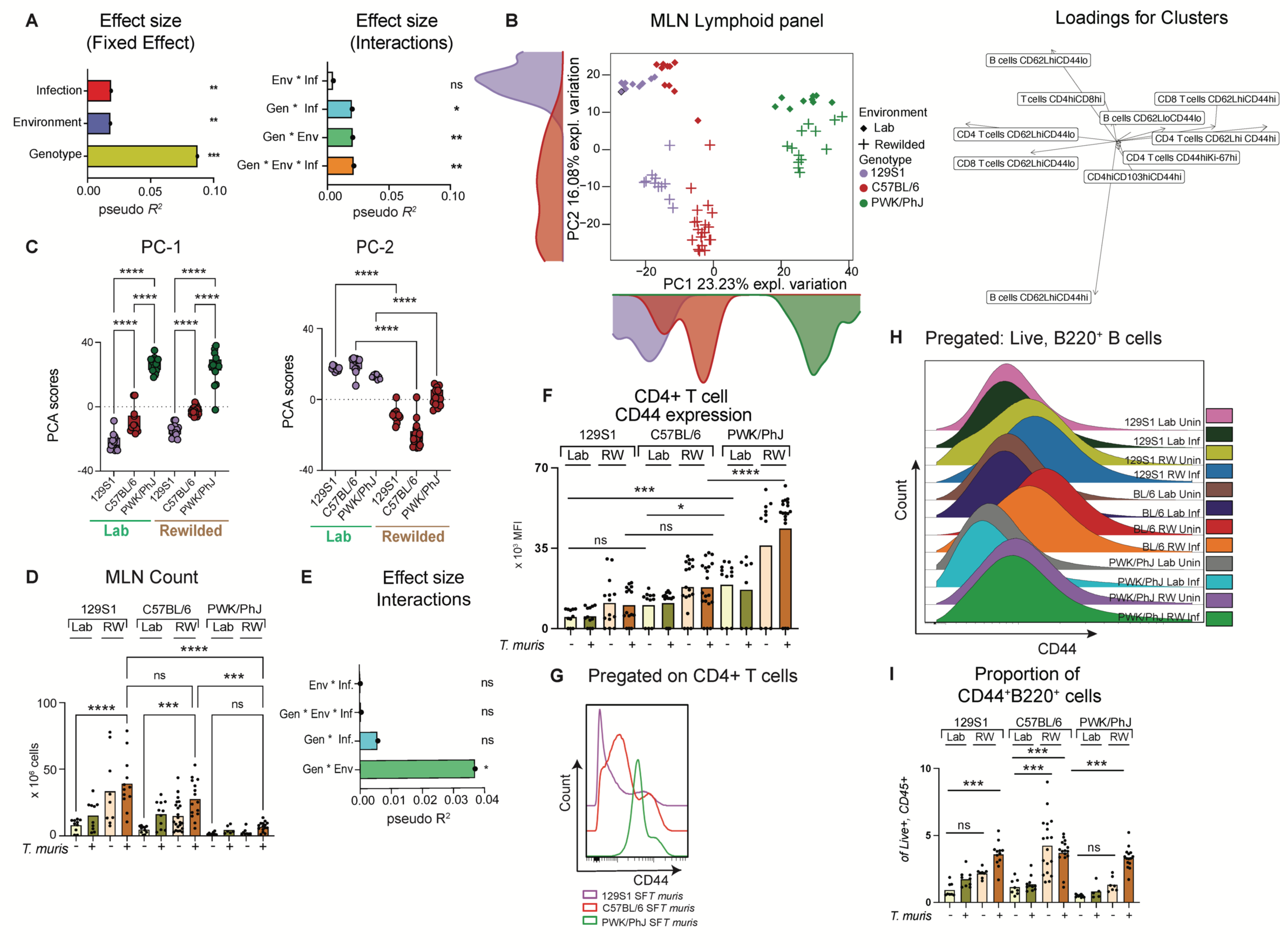
Interactions between Genotype, Environment and Infection contribute to variation in immune composition in murine MLNs. (**A**) Bar plots showing the pseudo *R^2^* measure of effect size of predictor variables and interactions as calculated by multivariate distance matrix regression analysis (MDMR). (**B**) PCA of immune cell clusters identified by unsupervised clustering in the MLN with the lymphoid panel and the loading factor of each population along the PCA. (**C**) Bar plot showing variance on PC1 and PC2 axis of PCA plots in (A). (**D**) MLN cell count from each mouse group and (**E**) pseudo *R^2^* measure of effect size of predictor variables and interactions as calculated by MDMR analysis based on MLN cell count (**F**) Bar plots depicting GMFI of CD44 on MLN CD4+ T cells. N = 5-15 mice per group, Block = 2. (**G**) Representative Histogram from Block 1 and 2 showing concatenated files from *T. muris* infected and rewilded mice of each mice strain (**H**) Representative Histogram showing concatenated files from different groups of mice in Block 2 with corresponding (**I**) Bar plots depicting proportion of B cells expressing CD44 on MLN cells. n = 6-18 mice per group, Block = 2. Statistical significance was determined by a based on MDMR analysis with R package for (**A**) and (**E**) or based on one-way ANOVA test between different groups with Graph-Pad Software for (**C**) and (**D**). For (**F**) and (**I**), one-way ANOVA test was used to test statistical significance between the different groups of interest. Data are displayed as mean ± SEM. *ns p>0.05; * p<0.05; ** p<0.01; *** p<0.001; **** p<0.0001*.

In addition to the independent effect of these factors, we also observed a G*E*I interaction in that for example, *T. muris* infection had a significant effect on cellular composition of the draining MLNs with increased proportion and sometimes abundance of B cells, especially in the 129S1 and the C57BL/6 strains, and especially following rewilding (Fig. S2D and S2E). Additionally, we observed that the morphology of the MLNs was quite different among mouse strains after rewilding, and this is reflected in the total cellular counts from the MLNs (Fig. 2D). Here, we observed that the PWK/PhJ mice had smaller lymph nodes that were not expanded in size compared to C57BL/6 and 129S1 mice after rewilding and *T. muris* infection, illustrating a Gen*Env interaction that could be statistically quantified by MDMR (Fig. 2E).

Further, we also observed based on the loading factors from the PCA analysis (Fig. 2B right) that the CD4 and B cell population in the MLN showed differential effect of the environment versus genetics in driving immune variation in the MLN. For example, as noted in the blood (Fig. 1E), we also found that in the MLN, expression of CD44 on CD4 T cells was majorly influenced by Genotype (Fig. 2F and Fig. 2G) with highest expression of CD44 on the PWK/PhJ mice across all environment while expression of CD44 on B cells, which usually depicts antigen experienced B cells (34) was predominantly influenced by Environment and exposure to *T. muris* parasites (Fig. 2H and I) with rewilded *T.muris* exposed mice of all genotypes having a higher percentage of CD44 expressing B cells than their counterparts in the vivarium (Fig. 2H, 2I and Fig. S2F).

These results indicate that Genotype and Infection (as well as Gen*Inf interactions) have a more significant effect on immune phenotypes in the draining lymph nodes than in the peripheral blood. Differences in lymph node size that are reflected in variation in lymph node cell numbers between different genotypes, as well as the residence of *T. muris* in the cecum, illustrates how the mesenteric lymph nodes provide a more complex picture of Gen*Env*Inf interactions than the peripheral blood.

### Genotype has the biggest effect on plasma and MLN stimulated cytokine levels

Cytokine response profiling is a common approach for immune phenotyping patients to characterize immune responses. Based on our previous study (23), we hypothesized that Genotype would have a larger effect on cytokine responses than on immune cell composition. Here, we measured both systemic circulating plasma cytokine levels in the plasma and cytokine production into the supernatants following microbial stimulation of the MLN cells. Indeed, as hypothesized, MDMR analysis of multiplex plasma cytokine data assessing systemic and circulating levels of IL-5, IL-17a, IL-22, IL-6, TNF-a and IFN-g (Table S8) showed that there are no statistically significant interactions among Genotype, Environment, and Infection (Fig. 3A, right); and the main effect of Genotype explained more variance than Environment (Fig. 3A, left). However, there is a strong effect of the different experimental blocks (Fig 3A), indicating that some unaccounted environmental or technical factors could also contribute to the variation. This Genotype effect on plasma cytokines (circulating levels of IL-5, IL-17a, IL-22, IL-6, TNF-a and IFN-g, Table S8) can be visualized on the PCA plot, where the C57BL/6 strain either in the lab or rewilded setting explained most of the difference on the PC1 axis (Fig 3B). Assessment of the loading factors revealed that the IFN-gamma levels were important in driving this variance (Fig 3C). Analysis of the cytokine data separately shows that the circulating IFN-γ levels were especially high in infected C57BL/6 mice in both lab and rewilded (Fig 3D).

**Fig. 3:**
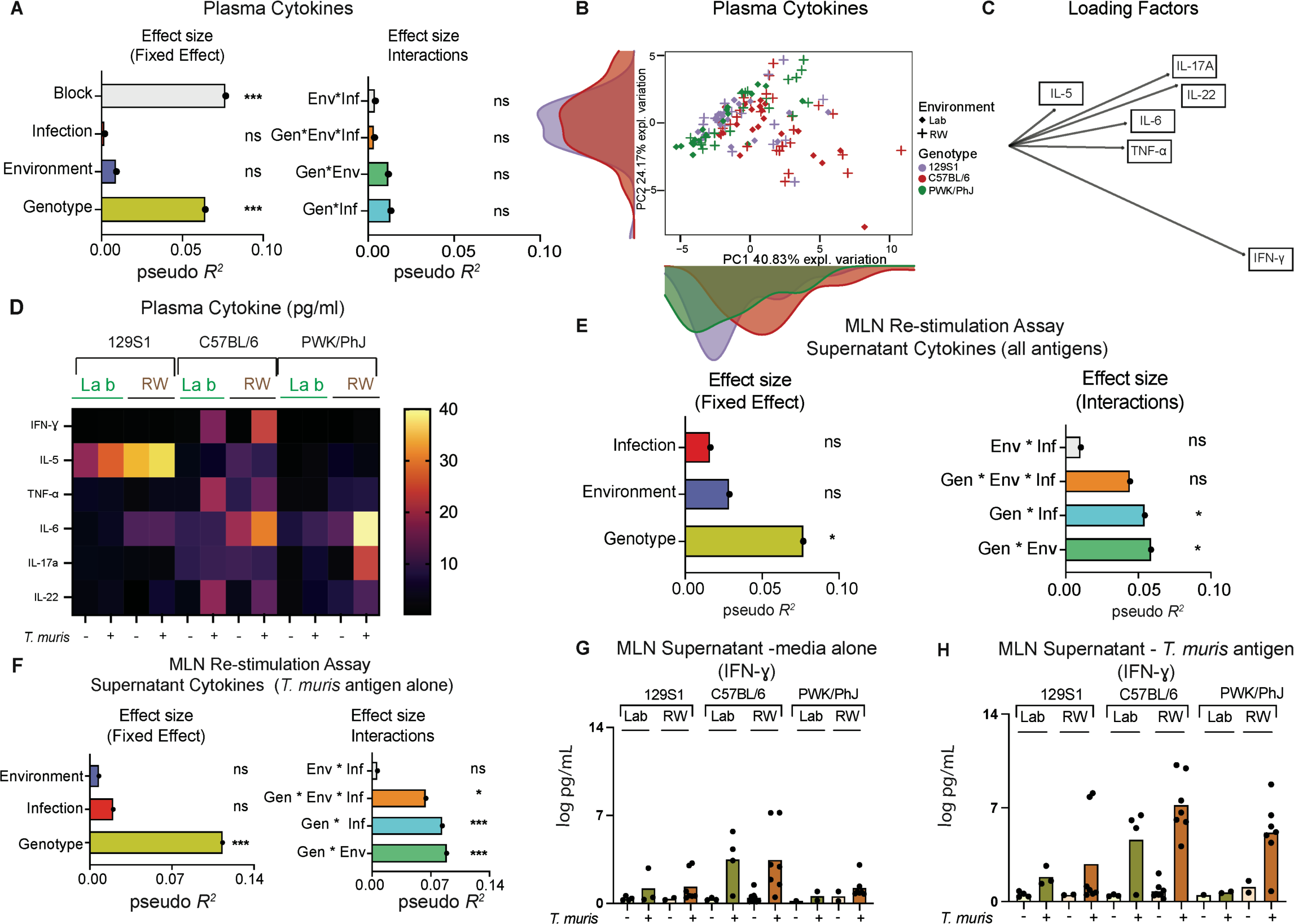
Genotype has a bigger effect on cytokine responses as a fixed predictor variable. (**A**) Bar plots showing the pseudo *R^2^* measure of effect size of predictor variables (left) and interactions (right) as calculated by multivariate distance matrix regression analysis (MDMR) from plasma cytokine data, Block 1 and Block 2 (**B**) PCA showing circulating plasma cytokine levels from individual mice and (**C**) their loading factors (**D**) Heatmap depicting levels of circulating cytokines in plasma (**E**) Bar plots showing the pseudo *R^2^*measure of effect size of predictor variables and interactions as calculated by multivariate distance matrix regression analysis (MDMR) from cytokine supernatant data of MLN cells stimulated with either CD3/CD28 beads, LPS, *Candida albicans*, *Clostridium perfringes*, *Bacteroides vulgatus* or *T. muris* antigens. (**F**) Bar plots showing the pseudo *R^2^* measure of effect size of predictor variables and interactions as calculated by multivariate distance matrix regression analysis (MDMR) from cytokine supernatant data of MLN cells stimulated with *T. muris* antigen. Barplot showing transformed IFN-γ cytokine levels in the supernatant for (**G**) controls as well as (**H**) following stimulation with *T. muris* antigen. Statistical significance was determined by a based on MDMR analysis with R package for (**A**), (**E**) and (**F**). Plasma cytokine data included samples from both Block 1 and Block 2 while MLN Cytokine supernatant data included samples from Block 2 alone due to technical problems with stimulation assays from Block 1. Data are displayed as mean ± SEM. ns *p>0.05; * p<0.05; ** p<0.01; *** p<0.001; **** p<0.0001*

When we characterized cytokine responses in the supernatant after *in vitro* stimulation of MLN cells, either with CD3/CD28 beads or with other microbial stimulants (LPS, *C. albicans*, *C. perfrigens*, *B. vulgatus* and *T. muris* antigens) (SFig 3, Table S9), MDMR analysis revealed Genotype having the biggest effect size on variation (Fig 3E, Left), which is consistent with the analysis of plasma cytokines. However, MDMR analysis of MLN cytokine responses also showed that the effect of Genotype on cytokine responses following stimulation of MLN cells with microbial antigens can be modulated by Environment and Infection (Gen*Env and Gen* Inf interactions) (Fig 3E, Right). When we focused our analysis on only cytokine responses to *Trichuris muris*, we confirmed that Genotype has the biggest effect on cytokine recall responses to *T. muris* antigen (Fig 3F, Left). However, this response also shows a significant effect of interactions between Gen*Env, Gen*Inf and Gen*Env*Inf (Fig 3F, Right). Analysis of the MLN supernatant cytokine data shows that consistent with the plasma cytokine data (Fig. 3D), production of IFN-γ from MLN cells also tends to be higher in C57BL/6 mice compared to the 129S1 strain of mice following *T. muris* infection in the lab or the rewilded environment (Fig. 3G and 3H).

Together, these results support our previous observations that genetics influence cytokine responses more strongly than the environment (23). However, here, we add evidence that the environment neither amplified nor eroded genetic effects on plasma cytokine levels, but that both environment and infection can modulate cytokine production in the MLN. Again, these results highlight the differences between analysis of the peripheral blood (the most accessible samples available in human studies), with analysis of draining lymph nodes that gather additional information on infections and the environment.

### Single Cell RNA Sequencing Analysis of mesenteric lymph node (MLN) cells emphasizes contribution of interactions between environment and genetics in immune variation

Single cell RNA sequencing (scRNA-seq) is an unbiased approach to profile immune phenotypes without pre-selection for analytes and markers of interest, and this approach can estimate both immune cell composition and provide insight into cytokine responses. Here, we used scRNA-seq to examine Gen*Env*Inf interactions on immune composition and cytokine responses in the MLN cells. Mesenteric lymph node cells (n=49,727) from individual mice (n=122) identified 23 major immune cell subsets visualized by UMAP (Fig. 4A). The cellular composition for each individual mice based on cluster membership with these 23 major immune cell subsets (Fig. S4A) is then used as the outcome variable for the MDMR analysis. In accordance with the cellular composition analysis with the flow cytometric data, MDMR analysis of the scRNAseq compositional dataset (Table S10) showed a significant effect of Genotype, Environment and Infection with *T. muris* in determining immune variation as fixed predictor variables in addition to a substantial block effect (Fig. 4B, above). Genotype and Environment (Gen*Env) interactions also explained significant variation in immune composition as assessed by scRNAseq (Fig. 4B, below). PCA analysis of the cellular composition from the single cell sequencing analysis (Fig S4A) of the different individual mice reveals contributions of Genotype and Environment to the variation among individual mice along the PC1 and PC2 axes (Fig. 4C). An example of how genotype effects can be modulated by environment (Gen*Env interaction), can be observed in the increase of Follicular B cells following rewilding that was especially heightened in C57BL/6 mice (Fig. 4D). In contrast, decrease in CD4 T cell abundance from rewilded naïve mice occurred for both 129S1 mice and C57BL/6 mice, but not PWK/PhJ mice (Fig. 4D and 4E). Overall, examination of cellular composition in the MLNs by scRNAseq resulted in a similar conclusion to spectral cytometry, in that Gen*Env interactions are particularly important. However, since we did not perform scRNAseq on the peripheral blood, we could not directly compare if Gen*Env interactions are more important in explaining variance in the cellular composition of MLNs than peripheral blood with this approach.

**Figure 4:**
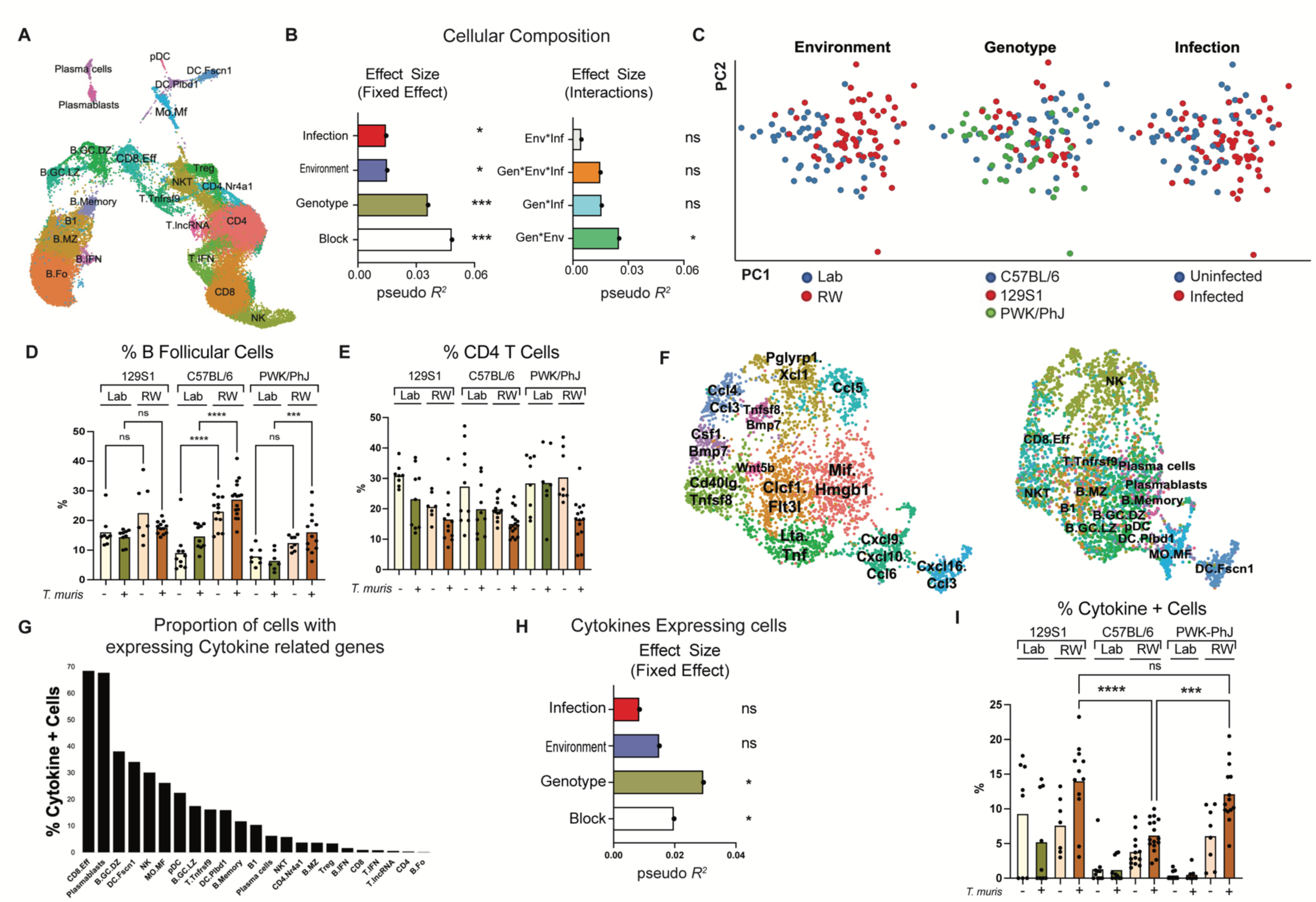
Single Cell Sequencing Analysis for assessing immune variation in cellular composition and cytokine profiles. (**A**) UMAP visualization of scRNASeq data identifying 23 major immune cell subset (**B**) Bar plot showing the pseudo *R^2^*measure of effect size of predictor variables and interactions as calculated by multivariate distance matrix regression analysis (MDMR) based on cellular composition of cells identified in (A) in each mouse (**C**) PCA of MLN cellular compositional data as determined by scRNAseq analysis Bar plots showing percentage of B Follicular cells (**D**) and CD4 T cells (**E**) based on the scRNA-Seq identified in (A). (**F**) Cytokine-expressing cell clusters. (**G**) Proportion of cells expressing cytokine-related genes of those identified in (F). (**H**) Bar plots showing the pseudo *R^2^* measure of effect size of predictor variables and interactions as calculated by multivariate distance matrix regression analysis (MDMR) based on data from proportion of cytokines expressing cells identified in (F). (**I**) Bar plot showing cytokine expressing cells identified in the (F). Statistical significance was determined by a based on MDMR analysis with R package for (**B**) and (**H**). Data are displayed as mean ± SEM. *ns p>0.05; * p<0.05; ** p<0.01; *** p<0.001; **** p<0.0001*.

To characterize the functional activity of the MLN cells, we used scRNAseq to identify the cells expressing cytokine related genes. Based on Gene Ontology, we extracted data for 232 genes defined to have molecular function in cytokine activity (GO:0005125), of which expression of 123 genes could be identified in the scRNAseq dataset (Table S11). Expression levels of these genes (n=123) were used to subset and re-cluster the MLN cells, and they were visualized based on expression of cytokine activity genes and their original cellular identity (Fig. 4F). Notably, CD4 T cells and Follicular B cells, which are the largest cellular populations in the overall dataset had the smallest percentage of cells that expressing cytokine genes (Fig. 4G), whereas CD8 effector cells, plasmablasts, and dark zone germinal center B cells, which are less abundant in the total population had higher proportions of cells expressing cytokine genes.

As described above, cells with cytokine activity were re-clustered based on their cytokine activity profiles (Fig 4F) and cluster membership with these cytokine activity subsets (Table S12) was then used as the outcome variable for the MDMR analysis. MDMR analysis showed that Genotype had a significant effect on variation in cells with cytokine activity (Fig. 4H), which is consistent with our previous work (23) and cytokine profiles described above. Also consistent with this analysis, other variables like Environment, Infection with *T. muris* (Fig. 4H) and Gen*Env or Gen*Env*Inf interactions (Fig. S4B) had no significant effect on variation in cellular cytokine activity as assessed by scRNA-Seq. The Genotype effect can be observed by plotting the percentage of MLN cells with cytokine activity for individual mice, with the 129S1 and PWK/PhJ mice having more cells expressing genes for cytokine activity than the C57BL/6 mice (Fig. 4I). PCA visualization of cellular composition based on cluster membership with cells of similar cytokine activity also showed distinct Genotype differences along the PC1 axis (Fig S4C).

Hence, an unbiased scRNAseq approach supports the conclusion that Genotype has the biggest effect on cytokine response heterogeneity, whereas cellular composition is more driven by interactions between Genotype and the Environment.

### Variation in *Trichuris muris* worm burden is determined by genetic, environmental and immunological factors

Ultimately, the question remains how the variance in these genetic, environmental and immunological factors influences susceptibility to subsequent infection. Therefore, we investigated predictors of worm burden (Fig. 5) and the contribution of genetics, environment and the different immunological factors to susceptibility to worm infection. Here, we observed that despite all 74 *T. muris*-exposed mice receiving approximately the same infectious dose (200 eggs), worm burden was negative-binomially distributed among exposed mice (Fig 5A). Analysis of worm burdens was done using Generalized Linear Models with a negative binomial error distribution. We found a significant Gen*Env for worm burden (Fig 5B, *p=0.04015*), whereby C57BL/6 mice harbored more worms than the other genotypes in the vivarium, but rewilding was associated with higher worm burdens in all genotypes. In other words, the relative susceptibility of the different host strains to *T. muris* depended upon environment (paralleling (35)). When we used logistic regression to analyze worm presence/absence at the experimental endpoint (reported as prevalence of infection among exposed mice in Fig. 5B), significant effects in the best model included only main effects of Genotype (*p=0.0001221*) and Environment (*p=0.0044835*), plus a significant effect of replicate experiment (*p=0.0329262*).

**Figure 5:**
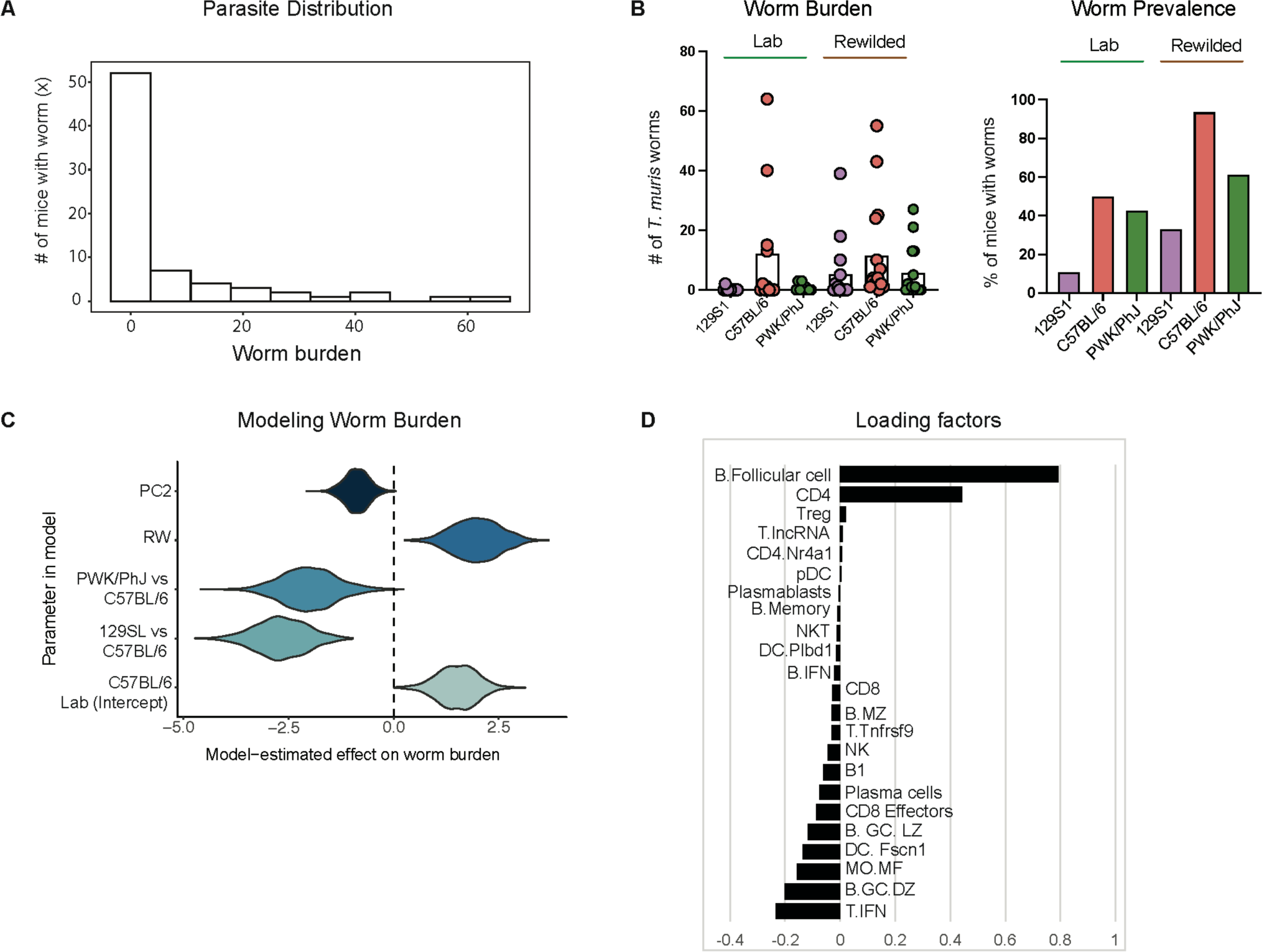
Genetics and Environmental factors predicts outcomes during exposure with *T. muris* parasite. Significant variation in worm burden among exposed mice, 3 weeks after inoculation with 200 eggs of *Trichuris muris* per host. (**A**) Worm burden (number of nematodes remaining in the caecum at that timepoint) followed a negative binomial distribution. (**B**) Worm burden depicted as Number of worms per mouse and Percentage of mice (Prevalence) still infected by worms. Each was predicted by a combination of Genetic and Environmental effects, including Gen*Env for worm burden (see text). (**C**) When we used PC2 from the scRNAseq data (Fig 3B) as an index of immune variation among individuals in our statistical models, we found that Gen*Env was no longer significant. Instead, the best model included main effects of host strain (C57BL/6 vs 129SL vs PWK/PhJ), environment (Lab vs RW), and PC2. The figure depicts 1000 model-estimated values for the effect of each predictor on worm burden. (**D**) Loading Factors for PC2 of the scRNAseq dataset.

Interestingly, when we included PC1 and PC2 values from the MLN scRNA-seq analysis (Fig. 5B) as summary measures of immune variation among individual mice, significant effects in the best model of worm burden (Fig. 5C) included only main effects of Genotype *(p=0.0003322)*, Environment (*p=0.0015615*), and PC2 scores *(p=0.0108213)*, which had a significant negative association with worm burden. Loading factors on the PC2 axis (Fig. 5D) indicated that the dearth of T cells with an interferon signature (*T.IFN*) may be a driver of the relationship between high PC2 scores and decreased worm burden. Furthermore, the fact that PC2 statistically outcompeted Gen*Env suggests that environment-dependent differences among genotypes in worm burdens may hinge on immune factors captured on PC2 (Fig5D). These results are consistent with T_H_1 responses being associated with increased susceptibility to helminth colonization (*36, 37*) and suggest that despite complexities in how immune phenotype is influenced by genetics and environment, once that immune phenotype emerges, established “rules” of infection susceptibility apply (as in (24)). The results also demonstrate that genetic, environmental, and individual immune variation is associated with varied infection burden.

## DISCUSSION

Our results support the hypothesis that the effect of even an extreme environmental shift on immune traits is modulated by genetics, and such modulations of phenotype through interactions between environment and genotype are an important source of variation in immune phenotypes. While we previously proposed that the immune cell composition for an individual is shaped by the environment (*23*), we find here that cellular composition in the peripheral blood is shaped by interactions between genetics and the environment and those in the mesenteric lymph node can shaped by interactions between genetics, environment and ongoing intestinal helminth infection. The role of genotype in shaping the structural lymph node non-immune cell environment as well as the lymphoid micro-environment as mediated by cytokines, chemokines and other stromal factors could explain the bigger effect of genotype in the lymph nodes compared to the peripheral blood. It is unsurprising that draining lymph nodes are more likely to detect and report immune phenotype changes that result from local tissue infections than the systemic circulatory immune cell populations. The complexity of Gen*Env*Inf interactions has important ramifications for the course of natural selection on the immune system, diversity in immune genotypes and efficacy of vaccine to pathogens. For example, because any given genotype may produce different immune responses in different environments, environment can alter the ability of individuals to resist and tolerate infections; furthermore natural selection operating on such variation is likely to generate divergent alleles and allele frequencies in different environments (*17*).

Quantification of such interactions is rare in immunological studies and is a valuable step forward in understanding the evolution and function of the immune system. The rewilding approach present a unique strategy for combining controlled mouse experiments with the advantages of tissue accessibility and homozygous mouse genetics, with increasing sophistication of multi-dimensional immune phenotyping analyses that have already been applied in human immunology. Based on these combinations, we do also identify traits for which main (i.e., non-interaction) effects are dominant. For instance, heterogeneity in cytokine responses shows a stronger influence of genetics, consistent with human studies (38). Furthermore, the Human Functional Genomics Project produced similar results to ours, that variation in T cell phenotypes is relatively more influenced by genetics, while B cell phenotypes are relatively more influenced by non-heritable environmental factors (1). It is thus likely that genetics and environment likewise contribute to human immune phenotype in synergistic ways. However, it is also possible that differences in interpreting genetic vs environmental contribution to human immune variation arise because of a focus on different immunological readouts (2, 3). We also found here that the contributions of Genotype, Environment and Infection could vary depending on tissue site analyzed, and since most human studies utilize peripheral blood, the effects of infection status depending on the infectious agent may appear less pronounced than if tissue samples were analyzed.

In comparing our results with human studies, it is also worth noting that human populations harbor greater heterozygosity and rarely undergo such dramatic environmental shifts as the laboratory mice being released into the rewilding environment. Longitudinal studies on travelers, refugees or immigrants may perhaps reveal similar alterations in immune phenotypes driven by environmental changes. Furthermore, the effect size of genotype versus environment on immune profile might also be influenced by other factors such as the age of the individual when the environmental change occurs. For example, a newborn might be more influenced by environmental factors immediately after birth where they are exposed to various environmental and microbial antigens as opposed to an adult (39–42). All the mice used in this study at the point of rewilding were all between 5-10 weeks of age, reflecting a young adult population in humans(43). Thus, perhaps releasing much younger mice or allowing sexual reproduction to occur in the rewilded environment might enhance the effects of environment; this remains a question for further investigation. Others have shown that interindividual variation tends to accumulate with age(44), thus, it will be interesting to evaluate the how other factors such as age can interact with environment and genetics to influence immune variation.

Conversely, using inbred strains of mice with homozygous alleles may represent an extreme test of genetic influences on immune variation. Compared to the inbred strains of mice that we have used in this study; most human genomes exist in a predominantly heterozygous state. Hence, despite these important differences between our approach and studies on human populations, there is surprising consistency between our findings here and previous conclusions from studies on human immune variation describing the differentially role of environment and genetics on B and T cell immune cell trait as well as the role of genetics in determine variation in cytokine responses (1, 38). However, our analysis and experimental design also present us an opportunity to assess the contribution of interaction to various immune traits and in different tissue sites. Together, this suggest that the rewilding experiments might be a bridge to study human immune response in murine models(21).

These experiments were performed in two experimental blocks over one summer. The effects of these two experimental blocks could either be technical variation in the analysis of samples, or biological variation resulting from changes to the environment over the two periods. Indeed, the block effects that we discuss in results such as the CBC/DIFF, cytokine levels and worm burden data could be driven by any number of technical or biotic factors that influence the temperature, diet and other commensals, which could subsequently influence immune responses in the mice, as well as differences in worm burden. With some assays (e.g. FACS and cytokine responses after MLN stimulations), the effects of experimental block is too great, so we analyzed the data generated as separate experiments instead of pooling data. Nonetheless, it is important to note that by including and removing the block effects statistically, we can quantify independent effects of Gen, Env and Inf, as well as the interactions between these variables while excluding experimental variation from the block effects. These findings further emphasize the role of environmental changes in influencing immune outcomes. A block effect could also explain some of the difference between the effect of rewilding on *T. muris* worm burden previously reported (24) and our findings within this study. In general, while some previous observations such as the significant expansion of the neutrophil pool across all genotype previously reported (26) and qualitative effect of rewilding on proportion of mice with *T. muris* (24) remain consistent across experimental time lapse, other outcomes such as number of worms per mice especially in the rewilded environment seems to be more dependent on the block effect and further emphasizing effect of the environment on immune outcomes.

Variation in adult worm burden of soil transmitted helminths between individuals typically follow a negative binomial distribution in humans (45) and wild animals (46). It is interesting to note that our quantification of larval stages in the process of being expelled also follow a negative binomial distribution. This raises the possibility that the negative binomial distribution in worm burdens in natural population may result from immune consequences of Gen*Env interactions as much as from differences in egg exposure. Hence, in addition to studies demonstrating that host genetics (47–49) and environmental factors (24) could influence susceptibility to helminth infection, we show here that interactions between genetics and environmental factors could also influence helminth infection outcomes. Variation in outcome to helminth infection is complex and maybe influenced by various other factors such as heterogeneity in parasite genetics factors (48–50), parasite dose and frequency of parasite exposure (51–54), host microbiome factors (55–57) as well as an individual’s infection history(4, 54) in nature. Importantly, natural helminth infection typically occurs from trickle infection of multiple small doses of egg exposures; therefore, the experimental system here of a high dose *T. muris* infection may not be representative a real-world exposure.

Critically, in this study, we also show that even systems such as this, where interactions between genetics and environment are important, the basic Th1 versus Th2 immunological mechanisms that govern susceptibility to *Trichuris muris* infection still predominate (24, 36). Thus, in addition, this study highlights a system where basic immunological mechanism that are established in the specific pathogen free facilities can be rigorously tested in a more naturalized system(21).

An interesting observation from these studies is the reduction in genetically driven immune phenotype differences in laboratory mice under rewilding conditions. This observation may be relevant to the different prevalence of inflammatory conditions across geographical locations – e.g., immune phenotypes may be more extreme in the absence of intensive microbial exposures and therefore have a greater impact in genetically susceptible individuals. One explanation of the hygiene hypothesis or the old friend’s hypothesis is that improved immune-regulatory responses through microbial exposure reduces the prevalence of inflammatory conditions (58–61). Our results also raise the possibility that increased microbial exposure may sometimes normalize or reduce the variation of immune phenotypes hence reducing the number of individuals with extreme immune responses. While the mechanism of this normalization is unclear, it is certainly possible that activation of immune-regulatory mechanisms through microbial exposure may normalize immune variation instead of simply reducing immune responses.

In conclusion, our results highlight how rewilding mice with controlled genetic backgrounds could be a bridge towards understanding immune variation between human individuals, and that quantification of the interactions at this interface may help elucidate the evolution of the immune system.

## Acknowledgements

We thank William Craigens, Christina Hansen and Felix Rozenberg for invaluable assistance in the field and at Stony Ford. In addition, we thank Mingming Zhao and Johnson Randall for help with setting up experiments in the lab and for help in maintaining the Joes’ Flow App Software respectively. We thank Dr. Elia Tait Wojno for sharing some *Trichuris muris* parasite eggs with us.

## Author contributions

Conceptualization: OO, AED, KC, ALG, PL. Methodology: OO, AED, NH, RSB, KK, YC, ALG, SCL, JD, OMP. Investigation: OO, AED, RB, NH, KK, KZ, YC, AM, SCL OMP, COSS, CH, PL. Data curation and analysis: OO, NH, AED, AG, SC. Writing – original draft: OO, AED, NH, ALG, PL. Writing – review and editing: OO, AED, KC, ALG, PL. Visualization: OO, NH, JC, AED, ALG. Supervision: SK, KC, ALG, PL. Funding acquisition: KC, ALG, PL.

## Data and materials availability

All data are available in the manuscript or the supplemental materials. Additional raw data, code and materials used in the analysis would be made publicly available at the point of publication.

## Materials and Methods Study Design

### Mice and Rewilding

C57BL/6J, 129S1/SV1mJ and PWK/PhJ mice were purchased from The Jackson Laboratory (Bar Harbor, ME) and were housed under specific pathogen–free conditions with *ad libitum* access to food and water. All mouse lines were then bred onsite in a specific pathogen free (SPF) facility at National Institute of Health (NIH). The resulting littermates from the multiple breeding pairs were shipped to Princeton University where they were randomly assigned to either remain in the institutional vivarium (lab mice) or released into the outdoor enclosures (rewilded mice) previously described and complementary manuscript (23–26). The protocols for releasing the laboratory mice into the outdoor enclosure facility were approved by Princeton IACUC.

25-30 mice of mixed strains and genotype, 129S1/SV1mJ, C57BL/6J and PWK/PhJ female mice were used for these experiments. For rewilding, 12-15 female mice of the different strains (129S1/SV1mJ, C57BL/6J and PWK/PhJ) were housed in different wedges in the enclosure for 5 weeks. Concomitantly, 10 mice of the different strains were left in the institutional vivarium (lab mice) where the temperature and humidity were maintained. Longworth traps baited with chow were used to catch the mice approximately at 2 weeks and 5 weeks after release(23). At 2 weeks after release, 8-10 mice of each genotype were trapped, blood and fresh stool was collected from the mice for longitudinal CBC analysis and microbiome analysis(25). At the same time 2 weeks following rewilding, some of these mice, vivarium controls and the rewilded mice, were infected with 200 *Trichuris muris* embryonated eggs by oral gavage. At approximately 5 weeks post-rewilding and 19-21 days post *T. muris* infection, mice were recovered for analysis and worm count. Blood was collected for Complete Blood Count with differentials and Flow Cytometry, MLN was harvested for flow cytometry and ceca were collected and the number of adult worms in each cecum were counted individually using an inverted microscope. For all analysis, samples that fail quality control and/or are under limit of detection are not included in downstream statistical analyses.

### Complete Blood Count Analysis

Blood samples (approx. 30-50μL) were collected from all mice at endpoint via cheek bleeds using a Medipoint Golden Rod Lancet Blade 4MM (Medipoint NC9922361) into a 1.3mL heparin coated tube (Sarstedt Inc, NC9574345). Blood samples were analyzed using the element HT5, Veterinary Hematology Analyzer (Heska).

### PBMC preparation and isolation

Heparinized Whole blood collected via the cheek bleeds were mixed with blood collected via cardiac puncture method. The combined blood samples were spun for 10 minutes at 1500 rpm and plasma was collected and stored at −80°C for further cytokine analysis. The cellular component re-suspended in PBS next underwent a density gradient separation process using the Lymphocyte Separation Media (LSM^TM^ MP Biomedicals, LLC, Irvine, CA) according to manufacturer’s instruction. Isolated PBMCs were washed twice in PBS and then used for downstream spectral cytometric analysis.

### Preparation of single cell suspensions from MLN

Single cell suspension from the mesenteric lymph nodes (MLNs) were prepared by mashing the tissues individually through a 70μm cell strainer and washed with RPMI. Cells were then washed with RPMI supplemented with 10% FCS. Live cell numbers were enumerated using the element HT5, Veterinary Hematology Analyzer (Heska).

### Spectral cytometry and Analysis

Single-cell suspensions prepared from the PBMCs, and MLN were washed twice with Flow cytometry buffer (FACs Buffer) and PBS prior to incubating with Live/Dead™ Fixable Blue (ThermoFisher) and Fc Block™ (clone KT1632; BD) for 10 minutes at room temperature. Cocktails of fluorescently conjugated antibodies (listed in Table 1 and Table 2) diluted in FACs Buffer and 10% Brilliant Stain Buffer (BD) were then added directly to cells and incubated for a further 30 minutes at room temperature. For the lymphoid panel, cells were next incubated in eBioscience™ Transcription Factor Fixation and Permeabilization solution (Invitrogen) for 12-18 hours at 4°C and stained with cocktails of fluorescently labeled antibodies against intracellular antigens diluted in Permeabilization Buffer (Invitrogen) for 1 hour at 4°C.

Spectral Unmixing was performed for each experiment using single-strained controls using UltraComp eBeads™ (Invitrogen). Dead cells and doublets were excluded from analysis. All samples were collected on an Aurora™ spectral cytometer (Cytek) and analyzed using the OMIQ software (https://www.omiq.ai/), data cleaning and scaling was done using algorithms like FlowCut (62, 63) within the OMIQ software. Sub-sampled cells including 10,000 Live, CD45+cells were re-clustered in an unsupervised version using the Joe’s Flow software, (Github: https://github.com/niaid/JoesFlow).

### MLN Cell Stimulation and Cytokine Profiling

Single cell suspension of MLN cells were reconstituted in RPMI at 2 × 10^6^ cells/mL, and 0.1 mL was cultured in 96-well microtiter plates that contained 10^7^ *cf.*u/mL UV-killed microbes, 10^5^ αCD3/CD28 beads (11456F), or Lipopolysaccharide (LPS; 100ng/mL) (L2630) or PBS control. The stimulated microbes and antigens included are as: *Bacteroides vulgatus* (ATCC 8482), *Candida albicans* (UC820), *Clostridium perfringens* (NCTC 10240), and *Trichuris muris* antigen. Supernatants were collected after 2 days and stored at −80°C. Concentrations of IL-5, IL-6, IL-22, IL-17A, IFN-γ, TNF-α, IL-2, IL-4, IL-10, IL-9 and IL-13 in supernatants were measured using a commercially available murine Th cytokine LEGENDplex assay (Biolegend) panel (Cat # 741044) according to the manufacturer’s instructions. Plasma concentrations of IL-5, IL-6, IL-22, IL-17A, IFN-γ, TNFα were measured also measured using the commercially available murine Th cytokine LEGENDplex assay (Biolegend) panel (Cat #741044) according to the manufacturer’s instructions. Cytokines levels that were lower than limit of detection across samples were excluded from further analysis.

### Single cell RNA sequencing

Single cell suspensions were obtained from MLN as described above. 2000 cells from each individual mouse (Block 1 (n = 51); Block 2 (n = 71)) were labelled with the Ab-Hash Tag Oligonucleoutides (HTO). These antibodies are a mix of anti-CD45 and anti-MHCI antibodies. TotalSeq^TM^-C antibodies are used with Single Cell 5′ kit. Pooled samples from each group were then loaded on a 10X Genomics Next GEM chip and single-cell GEMs were generated on a 10X Chromium Controller. Subsequent steps to generate cDNA and sequencing libraries were performed following 10X Genomics’ protocol. Libraries were pooled and sequenced using Illumina NovaSeq SP 100 cycle as per 10X sequencing recommendations.

The sequenced data were processed using Cell Ranger (version 6.0) to demultiplex the libraries. The reads were aligned to *Mus musculus* mm10 genomes to generate count tables that were further analyzed using Seurat (version 4.0). Sequencing data from the two blocks were integrated together prior to further downstream analysis. Data are displayed as uniform manifold approximation and projection (UMAP). The different cell subsets from each cluster were defined by the top 50 differentially expressed genes and identification using the SingleR sequencing pipeline(64). Cell types with different cytokine expression were identified based on expression of genes related to cytokine function using the Gene Ontology Browser. Seurat Analysis pipeline was used for comparisons between each of the different cell cluster of interest.

### Visualization

scRNASeq analysis data were visualized using Seurat (version 4.1.2) and R Studio (version 2022.07.1). Cartoons were created using BioRender.com.

### Quantification and Statistical analysis

In all cases principal component analysis was performed with in R v4.1.2. For cytokine data, log transformed data were used for generation of PCA plots and for MDMR analysis. Biplots were constructed by projecting the weighted averages of each input feature (immune cell phenotypes, cytokine level, cellular composition etc.,) along PC1, PC2 and/or PC3 derived from the biplot.pcoa function from the ape package as previously done in (23). Effect size measures were determined using the MDMR v0.5.1(65) package in R and interactions were tested using the mixed effect analysis in the MDMR package. A significant effect size is said to be present if the *p*-value was less than or equal to 0.05 (* *p*<0.05; ** *p*<0.01; *** *p*<0.001; **** *p*<0.0001).

Results in graphs and barplots are displayed as mean ± SEM using Prism version 7 (GraphPad Software, Inc.) except where mentioned. Statistical analysis was performed using GraphPad Prism software (v9). Right-skewed data were log or square root transformed. For analysis of relationship between scRNAseq cell composition and worm burden in Fig. 4, worm burden was modeled as following a negative binomial distribution. Predictor variables included in the regression model were mouse strain, mouse environment (i.e., laboratory or rewilded), and loading on PC2 from analysis of scRNAseq data. 1000 model-estimated coefficient values were then plotted for each predictor variable. In some cases, data were analyzed by One-Way ANOVA with Tukey’s post-when comparing three or more groups using GraphPad Prism software (v9). Experimental group was considered statistically significant if the fixed effect F test *p*-value was ≤0.05. Post hoc pairwise comparisons between experimental groups were made using Tukey’s honestly significant difference multiple-comparison test. A difference between experimental groups was taken to be significant if the *p*-value was less than or equal to 0.05 (* *p*<0.05; ** *p*<0.01; *** *p*<0.001; **** *p*<0.0001).

