## Supplemental Material for "Genetic and Environmental interactions contribute to immune variation in rewilded mice"

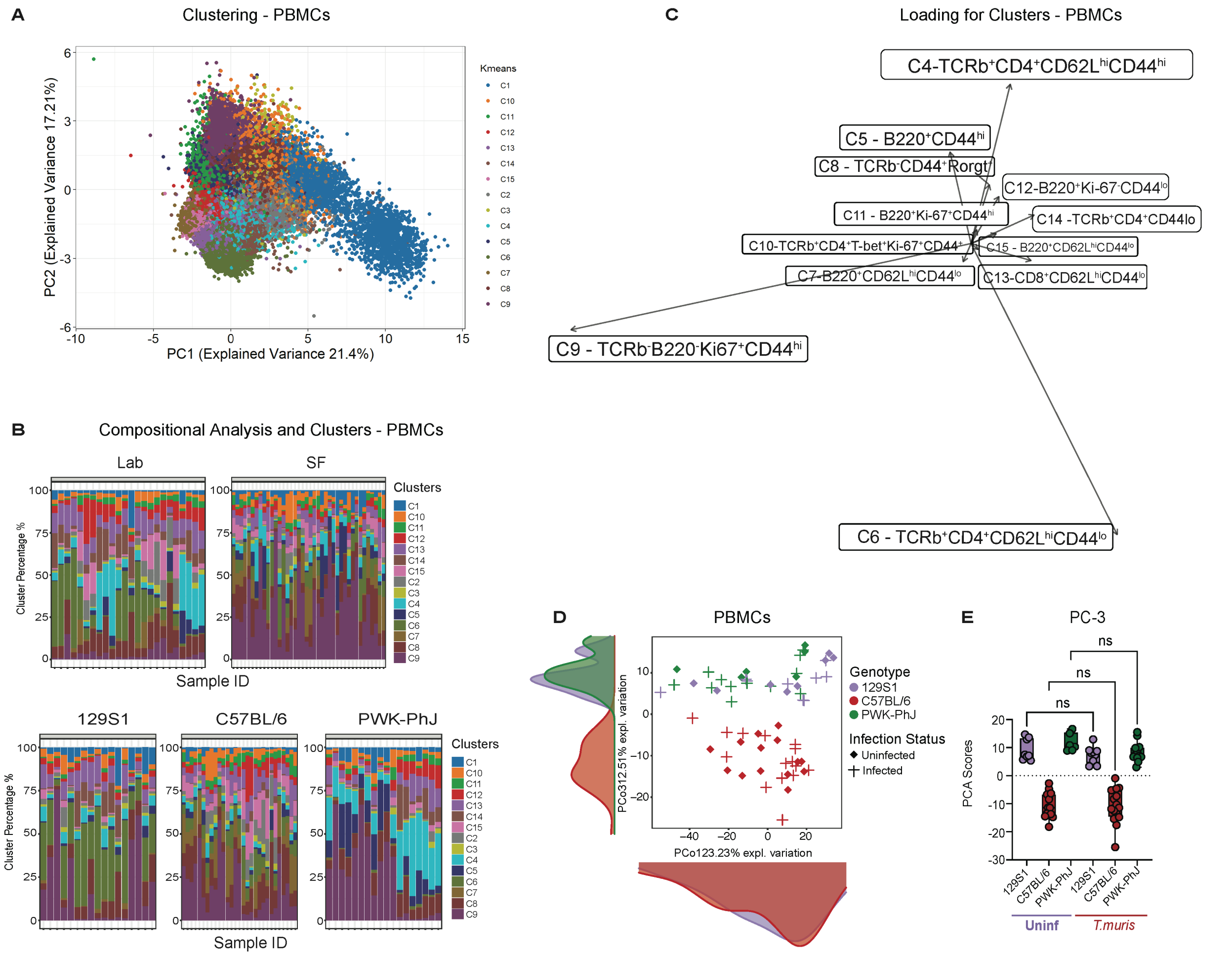

**SFig1: Unsupervised Clustering of PBMC cells.** (**A**) PCA Plot showing clusters generated following unsupervised clustering of PBMCs cells from all group of mice (**B**) Bar plot showing cluster percentage in different groups on a per mice basis. (**C**) the loading factors of immune clusters for PCA plot of PBMC cells showing PC1 and PC2 axis (**D**) PCA showing PC1 and PC3 axis of immune cell clusters identified by unsupervised clustering in the PBMCs cells and (**E**) the variance on PC1 vs PC3 axis.

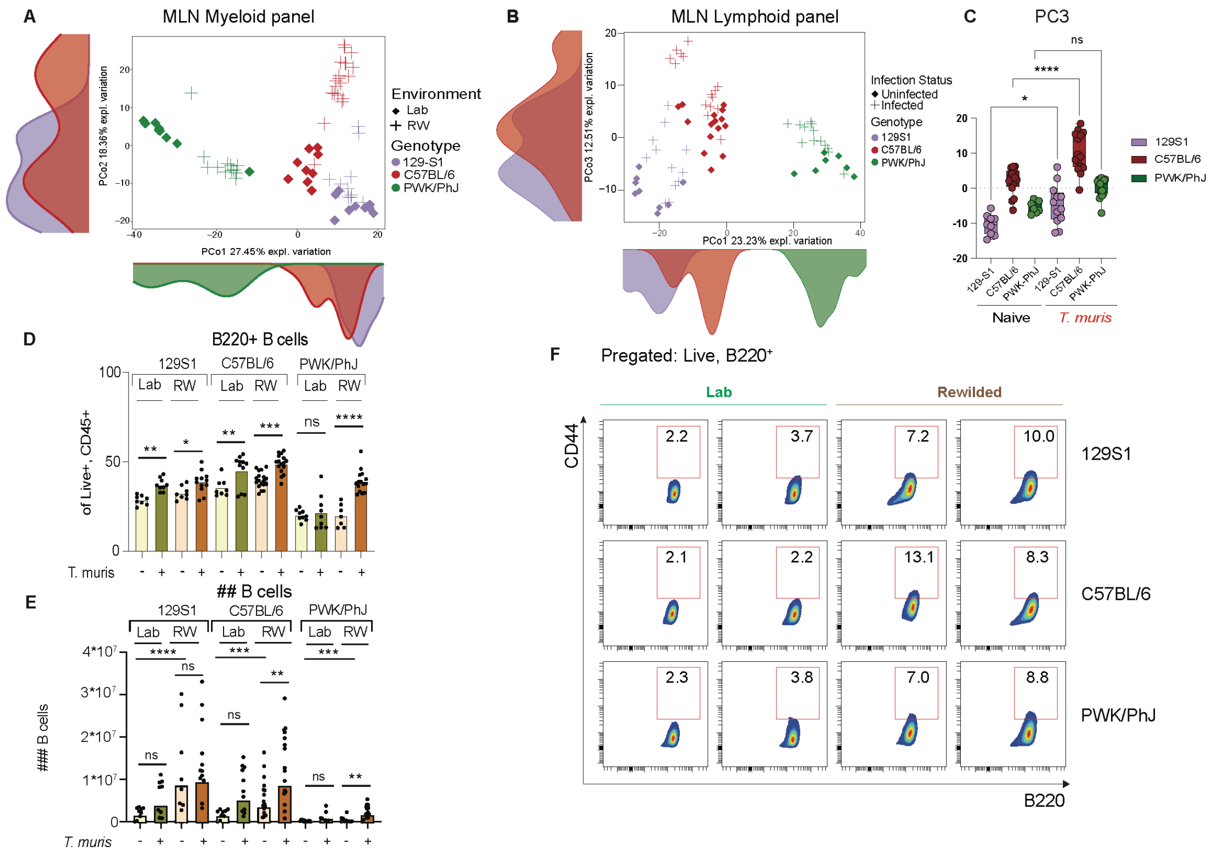


**SFig2: Interactions between Genotype, Environment and Infection determine immune composition in murine MLNs.** (**A**) PCA of immune cell clusters identified by unsupervised clustering in the MLN with the myeloid panel. (**B**) PCA of immune cell clusters identified by unsupervised clustering in the MLN with the lymphoid panel reflecting PC1 and PC3 with (**C**) Bar plot showing the variance on PC3 axis of PCA plots. Bar plots showing (**D**) frequencies and (**E**) numbers of B cells in the mesenteric lymph node (**F**) Representative FACS plots showing percentage of CD44 high B cells in the mesenteric lymph node. n = 6-18 mice per group, Block = 2. . Statistical significance was determined by one-way ANOVA test between different groups with Graph-Pad Software (**D**) and (**E**). For (**D**) and (**E**) direct comparison was done between groups of interest with one-way ANOVA test (within each genotype). Data are displayed as mean ± SEM. *ns p>0.05; * p<0.05; ** p<0.01; *** p<0.001; **** p<0.0001*.

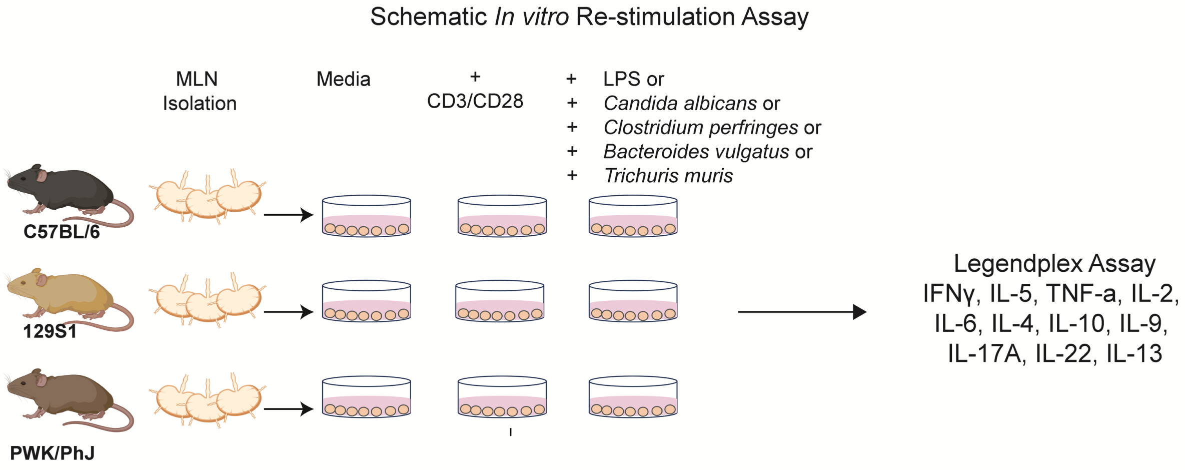


**SFig3: Schematic diagram of MLN *in-vitro* restimulation assay.** MLN cells from lab and rewilded of the different strains of mice were *ex-vivo* cultured with LPS, *C. albicans, C. perfringes, B. vulgatus, T. muris* or CD3/CD28 beads for 48 hours and supernatant was assayed for 11 cytokines IFN-γ, IL-5, TNF-α, IL-2, IL-6, IL-4, IL-10, IL-9, IL-17a, IL-22, IL-13.

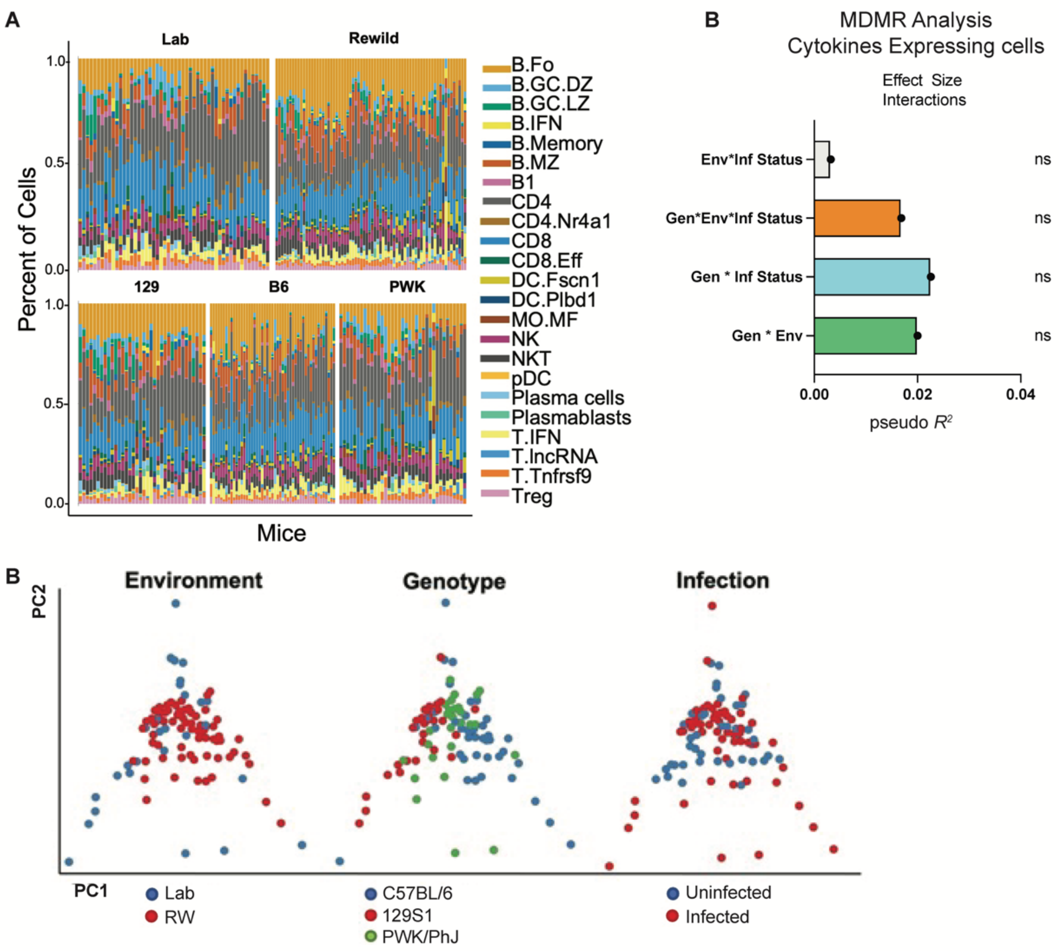


**SFig4: Single Cell Sequencing Analysis for assessing immune variation in cellular composition and cytokine profiles** (**A**) Proportion of cell types identified in Fig 3A on an individual mice basis (**B**) Bar plots showing the pseudo $R^2$ measure of effect size of interactions as calculated by multivariate distance matrix regression analysis (MDMR) for % of cell expressing cytokine genes (**C**) PCA of proportion of cytokine expressing cells as determined by scRNAseq analysis.

**Supplementary Materials**
**Materials and Methods**
**Table S1: Lymphoid Panel**

| REAGENT or RESOURCE | SOURCE | IDENTIFIER |
|---|---|---|
| Antibodies | | |
| Anti-mouse CD45 (30-F11) BUV395 | BD Biosciences | Cat#: 564279;<br>RRID: AB_2651134 |
| Rat Anti-mouse CD103 (M290) BUV496 | BD Biosciences | Cat#: 741083;<br>RRID: AB_2870687 |
| Rat Anti-mouse CD62L (MEL-14) BUV563 | BD Biosciences | Cat#: 741230;<br>RRID: AB_2870784 |
| Hamster Anti-mouse TCR-B chain (H57-597) BUV737 | BD Biosciences | Cat#: 612821 |
| Rat Anti-Mouse CD44 (IM7) BUV805 | BD Biosciences | Cat#: 741921<br>RRID: AB_2871234 |
| Rat Anti-Mouse NKp46 CD335 (29A1.4), BV480 | BD Biosciences | Cat#: 746264<br>RRID: AB_2743596 |
| Anti-Mouse NK-1.1 (PK136), BUV661 | BD Biosciences | Cat#:741477<br>RRID: AB_2870942 |
| Hamster Anti-Mouse TCR β Chain, (H57-597), BUV737 | BD Biosciences | Cat#: 612821 |
| Anti-mouse CD8β (H35-17.2) BV510 | BioLegend | Cat#:103278<br>RRID: AB_2860603 |
| Anti-mouse CD4 (RM4-5) BV570 | BioLegend | Cat#:100542<br>RRID: AB_2563051 |
| Hamster Anti-Mouse KLRG1 (2F1), BV750 | BD Biosciences | Cat#: 746972;<br>AB_2871754 |
| Anti-mouse TCRγδ (eBio-GL3) PECy5 | Thermofisher | Cat#**: 15-5711-82**<br>RRID: AB_468804 |
| Anti-mouse CD45R/B220 (RA3-6B2) APC/Fire810 | BioLegend | Cat#: 103278<br>RRID: AB_2860603 |
| Ant-mouse Tbet (4B10) BV421 | Biolegend | Cat#": 644816<br>RRID: AB_10959653 |

| | | |
|---|---|---|
| Ki-67 Monoclonal Antibody (SolA15) AF532 | Thermofisher | Cat#: 58-5698-82<br>RRID: AB_2802365 |
| Anti-mouse FoxP3 (FJK-16) AF700 | BioLegend | Cat#: 320014<br>RRID: AB_439750 |
| Anti-mouse RorgT (Q31-378) PE-CF594 | BD Biosciences | Cat#: 56284<br>RRID: AB_2651150 |
| Gata-3 Monoclonal Antibody (TWAJ), Alexa Fluor™ 488 | Thermofisher | Cat#: 53-9966-42;<br>RRID: AB_2574493 |
| Hamster Anti-Mouse CD69 (H1.2F3) BV650 | BD Biosciences | Cat#: 740460;<br>RRID: AB_2740186 |
| Anti-Mouse CD279 (PD-1) (29F.1A12) BV421 | BioLegend | Cat#:135218;<br>RRID: AB_2561447 |
| Anti-mouse CD16/32 Fc Block™ (KT1632) | BD Biosciences | Cat#: **MA5-18012**<br>RRID: AB_2539396 |

**Table S2: Myeloid Panel**

| **REAGENT or RESOURCE** | **SOURCE** | **IDENTIFIER** |
|---|---|---|
| **Antibodies** | | |
| Anti-mouse CD45 (30-F11) BUV395 | BD Biosciences | Cat#: 564279;<br>RRID: AB_2651134 |
| Rat Anti-mouse CD43 (S7) BUV563 | BD Biosciences | Cat#: 741238;<br>RRID: AB_2870790 |
| Anti-mouse CD11b (M1/70) BUV615 | BD Biosciences | Cat#: 751140<br>RRID: AB_2875166 |
| Anti-mouse TCRβ (H57-597) BUV661 | BD Biosciences | Cat#: 749914<br>RRID: AB_2874153 |
| Anti-mouse CD44 (IM7) BUV805 | BD Biosciences | Cat#: 741921<br>RRID: AB_2871234 |
| Anti-Mouse CD279 (PD-1) (29F.1A12) BV421 | BioLegend | Cat#: 135218;<br>RRID: AB_2561447 |
| MERTK Monoclonal Antibody (DS5MMER), Super Bright 436 | Thermofisher | Cat#: 62-5751-82<br>RRID: AB_2688137 |
| Anti-mouse CD49b (DX5), Pacific Blue | Biolegend | Cat #: 108918;<br>RRID: AB_2265144 |

| | | |
|---|---|---|
| Rat Anti-mouse F4/80 Anti-Mouse (T45-2342) F4/80 | BD Bioscience | Cat #: 565635<br>RRID: AB_2739313 |
| Hamster Anti-Mouse CD27 (CD27) | BD Bioscience | Cat #: 563605;<br>RRID: AB_2738310 |
| Anti-mouse CD8α (5H10) Pacific Orange | Thermofisher | Cat#: MCD0830<br>RRID: AB_10376311 |
| Anti-mouse CD4 (RM4-5) Qdot800 | Thermofisher | Cat#: **Q22165**<br>RRID:AB_2556521 |
| Anti-mouse CD64 (X54-5/7.1) PECy7 | BioLegend | Cat#: 139323<br>RRID: AB_2629778 |
| Anti-human CD278 (ICOS) (C398.4A) BV510 | Biolegend | Cat #: 313525<br>RRID: AB_2562642 |
| Anti-Mouse CD11c (N418) BV711 | Biolegend | Cat#: 117349<br>RRID: AB_2563905 |
| Hamster Anti-Mouse CD183 (CXCR3-173) BV750 | BD Biosciences | Cat#: 747298<br>RRID: AB_2872012 |
| Anti-Mouse CX3CR1 (SA011F11) BV785 | Biolegend | Cat#: 149029<br>RRID: AB_2565938 |
| Rat Anti-mouse Siglec F (E50-2440) BB515 | BD Bioscience | Cat#:564514<br>RRID: AB_2738833 |
| Anti-mouse TCRγδ (eBio-GL3) PECy5 | Thermofisher | Cat**#: 15-5711-82**<br>RRID: AB_468804 |
| Anti-mouse CD301b/MGL2 (URA-1) PerCPCy5.5 | BioLegend | Cat#:146810<br>RRID: AB_2563391 |
| Anti-mouse CD273/PDL2 (B7-DC) PE | BioLegend | Cat#:115565<br>RRID: AB_2819827 |
| Anti-mouse CD45R/B220 (RA3-6B2) APC/Fire810 | BioLegend | Cat#: 103278<br>RRID: AB_2860603 |
| Rat Anti-mouse CD62L (MEL-14) BUV563 | BD Biosciences | Cat#: 741230;<br>RRID: AB_2870784 |
| Anti-mouse CD19 Antibody (6D5) Spark Blue 550 | Biolegend | Cat#: 115566<br>RRID: AB_2832389 |
| Mouse CXCR5 (614641) Alexa Fluor® 488 | R&D | Cat#: FAB6198G-100UG |
| Anti-mouse CD69 (H1.2F3) PE-Dazzle 594 | Biolegend | Cat#: 104535<br>RRID: AB_2565583 |
| Anti-mouse CD25 (PC61.5), PE-Cy5 | Thermofisher | Cat#: 15-0251-82<br>RRID: AB_468733 |

| | | |
|---|---|---|
| B7-2/CD86 (BU63) PE-Cy5.5 | Novus Biological | Cat#: NBP2-34569PECY55 |
| Anti-mouse Tim-4 (RMT4-54) PECy7 | Biolegend | Cat#: 130010<br>RRID: AB_2565719 |
| Anti-mouse ST2 (DIH4) APC | Biolegend | Cat#: 146606<br>RRID: AB_2728175 |
| Anti-mouse CD206 (MMR) AF-647 | Biolegend | Cat#: 141712<br>RRID: AB_10900420 |
| Anti-mouse Ly-6c (HK1.4), PerCP | Biolegend | Cat#: 128028<br>RRID: AB_10897805 |
| Anti-mouse Ly-6G (1A8), Spark NIR 685 | Biolegend | Cat#: 127666<br>RRID: AB_2876454 |
| Anti-mouse CD103, APC-R700 | BD Biosciences | Cat#: 565529<br>RRID: AB_2739282 |
| Anti-mouse KLRG1 (2F1/KLRG1), APC-Cyanine 7 | Biolegend | Cat#: 138436,<br>RRID: AB_2566554 |
| Anti-mouse CD16/32 Fc Block™ (KT1632) | BD Biosciences | Cat#: **MA5-18012**<br>RRID: AB_2539396 |

**Table S3: HashTag Oligonucleotides (HTOs)**

| REAGENT or RESOURCE | SOURCE | IDENTIFIER |
|---|---|---|
| **Antibodies** | | |
| TotalSeq™ -C0301 anti-mouse Hashtag 1 Antibody (M1/42; 30-F11) | BioLegend | 155861 |
| TotalSeq™ -C0302 anti-mouse Hashtag 2 Antibody (M1/42; 30-F11) | BioLegend | 155863 |
| TotalSeq™ -C0303 anti-mouse Hashtag 3 Antibody (M1/42; 30-F11) | BioLegend | 155865 |
| TotalSeq™ -C0304 anti-mouse Hashtag 4 Antibody (M1/42; 30-F11) | BioLegend | 155867 |
| TotalSeq™ -C0305 anti-mouse Hashtag 5 Antibody (M1/42; 30-F11) | BioLegend | 155869 |
| TotalSeq™ -C0306 anti-mouse Hashtag 6 Antibody (M1/42; 30-F11) | BioLegend | 155871 |
| TotalSeq™ -C0307 anti-mouse Hashtag 7 Antibody (M1/42; 30-F11) | BioLegend | 155873 |
| TotalSeq™ -C0308 anti-mouse Hashtag 8 Antibody (M1/42; 30-F11) | BioLegend | 155875 |
| TotalSeq™ -C0309 anti-mouse Hashtag 9 Antibody (M1/42; 30-F11) | BioLegend | 155877 |
| TotalSeq™ -C0310 anti-mouse Hashtag 10 Antibody | BioLegend | 155879 |

| **Bacterial and Antigens, Ex-vivo stimulations** | | |
|---|---|---|
| *Lipopolysaccride* | | |
| *Candida albicans* | ATCC | UC820 |
| *Clostridium perfringens* | NCNC | NCTC 10240 |
| *Bacteroides vulgatus* | ATCC | ATCC 8482 |
| *T. muris excretory secretory products (TES)* | Generated in-house | (62, 63) |

| | | |
|---|---|---|
| Mouse T-Activator CD3/CD28 for T cell expansion and activation | Thermofisher | 11456D |

| **Experimental Models: Organism/Strains** | | |
|---|---|---|
| C57BL/6J | The Jackson Laboratory and bred in house | JAX: 000664 |
| 129S1/SV1MJ | The Jackson Laboratory | JAX: 002448 |
| PWK/PhJ | The Jackson Laboratory | JAX: 003715 |
| *Trichuris muris* | Tait Wojno Lab | N/A |

| **Critical Commercial Assays and Materials** | | |
|---|---|---|
| Live/Dead Fixable Dead Cell Stain Kits | Invitrogen | Cat#: L23105 |
| Brilliant Stain Buffer Plus | BD Biosciences | Cat#: 566365 |
| LEGENDplex™ MU TH Cytokine panel (12-plex) w/FP V03 | Biolegend | Cat#: 741043 |
| Element HT5, Diluent | Heska | Cat#:5221 |
| Element HT5, Probe Cleanser | Heska | Cat#: 5228-1 |
| Element HT5, Diff Lyse Solution | Heska | Cat#: 5223 |

| | | |
|---|---|---|
| Element HT5, LH Lyse Solution | Heska | Cat#: 5225 |
| QIAsymphony PowerFecal Pro DNA Kit | Qiagen | Cat#: 938036 |
| Chromium Next GEM Chip G Single Cell Kit | 10X Genomics | Cat#: 1000120 |
| Chromium Next GEM Single Cell 5' Library & Gel Bead Kit v1.1, | 10X Genomics | Cat#: 1000165 |
| Chromium Single Cell 5' Library Construction Kit | 10X Genomics | Cat#: 1000020 |
| Single Index Kit T Set A, 96 rxns | 10X Genomics | Cat#: 1000213 |

| **Software and Algorithms** | | |
|---|---|---|
| SpectroFlo® | Cytek | N/A |
| Joe's Flow | Devlin *et al* in preparation | https://github.com/niaid/JoesFlow/ |
| Prism v9 | GraphPad | N/A |
| Seurat v3 | 43 | https://satijalab.org/seurat/ |
| R v4.1.2 | R Studio, PBC | https://www.r-project.org/ |
| OMIQ | OMIQ | https://www.omiq.ai |
| JMP v16 | SAS | N/A |
| Illustrator CC | Adobe | https://www.adobe.com/products/illustrator.html |
| Algorithms: Effect size measures | MDMR v0.5.1 package | https://link.springer.com/article/10.1007/s11336-016-9527-8 |
